# The microtubule-severing enzyme spastin regulates spindle dynamics to promote chromosome segregation in *Trypanosoma brucei*

**DOI:** 10.1101/2025.01.03.631140

**Authors:** Thiago Souza Onofre, Qing Zhou, Ziyin Li

## Abstract

Microtubule-severing enzymes play essential roles in regulating diverse cellular processes, including mitosis and cytokinesis, by modulating microtubule dynamics. In the early branching protozoan parasite *Trypanosoma brucei*, microtubule-severing enzymes are involved in cytokinesis and flagellum length control during different life cycle stages, but none of them have been found to regulate mitosis in any life cycle form. Here, we report the biochemical and functional characterization of the microtubule-severing enzyme spastin in the procyclic form of *T. brucei*. We demonstrate that spastin catalyzes microtubule severing *in vitro* and ectopic overexpression of spastin disrupts spindle microtubules *in vivo* in trypanosome cells, leading to defective chromosome segregation. Knockdown of spastin impairs spindle integrity and disrupts chromosome alignment in metaphase and chromosome segregation in anaphase. We further show that the function of spastin requires the catalytic AAA-ATPase domain, the microtubule-binding domain, and the microtubule interacting and trafficking domain, and that the association of spastin with spindle depends on the microtubule-binding domain. Together, these results uncover an essential role for spastin in chromosome segregation by regulating spindle dynamics in this unicellular eukaryote.

## Introduction

*Trypanosoma brucei* is an early branching protozoan causing sleeping sickness in humans and nagana in cattle in sub-Saharan Africa. This unicellular parasite has a complex life cycle by alternating between the insect vector and the mammalian host, within which it proliferates through binary fission along its longitudinal axis using mechanisms that are distinct from the host cells. Trypanosomes undergo a closed mitosis without breaking down the nuclear envelope, and assembles an intra-nuclear mitotic spindle to separate the duplicated chromosomes (1). In eukaryotes, the mitotic spindle is a highly dynamic molecular machine composed of microtubules, motors and non-motor proteins, which modulate microtubule dynamics and orchestrate physical forces to separate the sister chromatids (2). Assembly of a bipolar spindle depends on the centrosome/spindle pole body, which serves as the microtubule-organizing center (MTOC), and the spindle pole-localized γ-tubulin ring complex (γ-TuRC), which serves as the microtubule nucleation machinery (2). In *T. brucei*, however, a canonical centriole structure is not detectable at the spindle pole, but a ring-like structure, from which spindle microtubules emanate, is present at the spindle pole (1), suggestive a likely MTOC. Intriguingly, the γ-TuRC in *T. brucei* contains an unusual subunit composition and plays no essential roles in spindle assembly (3,4), despite localizing to the spindle poles (5), indicating a γ-TuRC-independent mechanism in spindle microtubule nucleation. Moreover, close homologs of several essential mitotic regulators in yeast and humans, including the spindle motor protein BimC and the kinetochore motor protein CENP-E, are missing from the genome (6). Thus, trypanosomes likely employ distinct control mechanisms to regulate spindle assembly and chromosome segregation, necessitating the in-depth exploration of the mechanism of mitosis regulation in *T. brucei*.

The nuclear genome of *T. brucei* consists of 11 pairs of homologous megabase chromosomes (1-6Mb in size), which have regional centromeres, and >100 sub-megabase chromosomes, including ∼100 mini-chromosomes of 50-150kb in size and 3-5 intermediate-chromosomes of 200-700kb in size, which do not have canonical centromeres (7). Mini-chromosomes segregate via spindle microtubule-dependent mechanisms, likely by attaching to the central non-kinetochore microtubules, and they have segregation kinetics different from the megabase chromosomes (8). The mitotic spindle in *T. brucei* possesses two types of clearly identifiable microtubules, kinetochore microtubules and pole-to-pole microtubules or non-kinetochore microtubules (9), and multiple bundles of up to 20 microtubules are detectable in a mitotic cell (8). Astral microtubules that extend from the spindle poles are, however, not present as in animal cells, and the spindle poles appear to be anchored on the nuclear envelope (8). The kinetochore on the megabase chromosomes has a characteristic trilaminar structure (8), and is composed of ∼20-30 proteins, most of which are unique to trypanosomes (10–13), suggestive of an unusual molecular composition. Notably, *T. brucei* lacks the homologs of the conserved spindle-assembly checkpoint machinery that monitors kinetochore-microtubule attachment errors; thus, it is unclear how this parasite monitors and corrects such errors to maintain genome stability.

Chromosome segregation occurs in anaphase that consists of anaphase A, during which chromosomes move toward the spindle poles via depolymerization of kinetochore microtubules, and anaphase B, during which spindle poles move apart via elongation of the spindle (14–16). Depolymerization of kinetochore microtubules at the plus ends generates chromosome movement termed “Pacman”, and depolymerization of kinetochore microtubules at the minus ends generates chromosome movement termed “poleward flux” (17). The kinesin-13-family microtubule depolymerase and the microtubule-severing enzyme katanin contribute to chromosome movement in anaphase A through the Pacman mechanism (18,19), whereas the microtubule-severing enzymes fidgetin and spastin contribute to chromosome movement in anaphase A through the poleward-flux mechanism (19). In *T. brucei*, both the kinesin-13-family microtubule depolymerases and the microtubule-severing enzymes, katanin, spastin, and fidgetin, are expressed. Kif13-1/KIN13-1 regulates spindle dynamics and chromosome segregation (20,21), although its mode of action remains unclear. In the bloodstream form, all four katanin proteins, KAT60a, KAT60b, KAT60c, and KAT80, are involved in cytokinesis (22), whereas in the procyclic (insect) form, the KAT60a-KAT80 complex is required for cytokinesis (23) and KAT60b is involved in flagellum length control (24). Spastin is required for cytokinesis in the bloodstream form (22), and fidgetin is nonessential in both the procyclic and the bloodstream forms (22), despite localizing to the nucleus and the spindle (25). None of these microtubule-severing enzymes have previously been demonstrated to play a role in mitosis in *T. brucei*.

In this work, we characterized the function of spastin in the procyclic form of *T. brucei* and showed that it associates with the mitotic spindle and is required for faithful chromosome segregation. We also demonstrated its *in vitro* microtubule-severing activity and identified its structural motifs required for function. These results uncovered important roles of spastin in regulating spindle dynamics to promote chromosome segregation in this early divergent microbial eukaryote.

## Results

### *T. brucei* spastin severs microtubules *in vitro*

We performed bioinformatic analysis and structural modeling/prediction of the *T. brucei* spastin homolog, TbSPA, and found that it contains an unusual N-terminal extension of a BRCT (BRCA1 C-terminal) domain, a protein-protein interaction module found in numerous proteins in prokaryotes and eukaryotes (26), when compared with human spastin (Fig. 1A, B). The BRCT domain adopts a structure with three α-helices surrounding a central β-sheet (Fig. 1B), and in human BRCT-containing proteins this domain binds phosphopeptides and mediates signal transduction (27). TbSPA contains a conserved AAA domain (<u>A</u>TPase <u>A</u>ssociated with diverse cellular <u>A</u>ctivities) at its C-terminus and multiple motifs found in human spastin, the hydrophobic domain (HD), the microtubule interacting and transport (MIT) domain, and the microtubule-binding domain (MBD), in the middle region (Fig. 1A, B). The AAA domain in TbSPA is predicted to adopt a nucleotide-binding domain (NBD) comprised of α-helices/β-sheets and a helix-bundle domain (HBD) comprised of several α-helices (Fig. 1B), typical of the AAA domain. In human spastin, the HD and the MIT domains are involved in binding to different proteins and are required for different cellular activities; thus, these two domains appear to target spastin to different subcellular locations. The MBD domain binds to microtubules in an ATP-independent manner and is required for the severing activity, which is ATP dependent and is mediated by the AAA domain (28).

**Figure 1.**
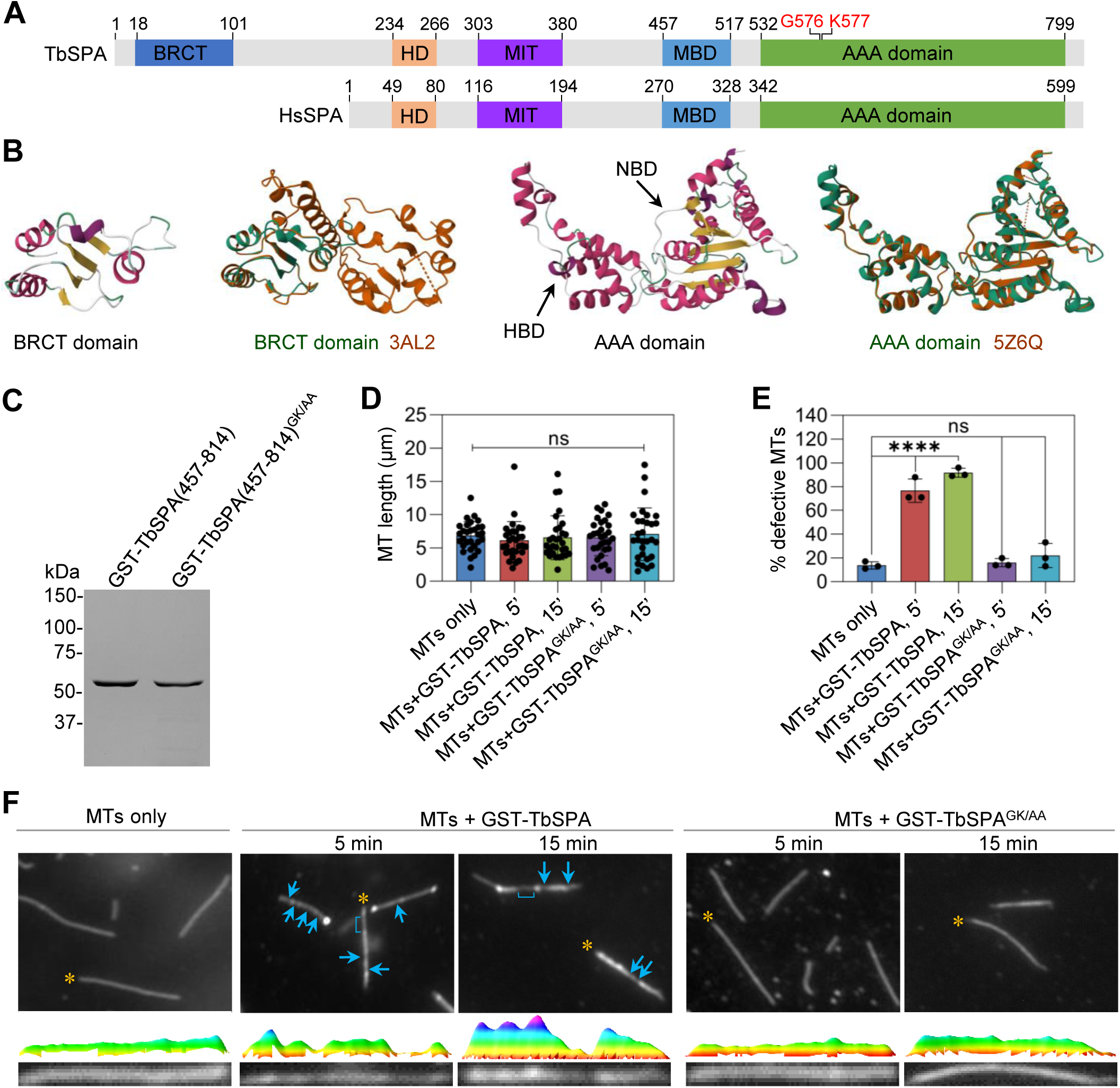
Trypanosome spastin catalyzes microtubule severing *in vitro*. **(A)**. Schematic drawing of T. brucei spastin (TbSPA) structural domains and comparison with human spastin (HsSPA). BRCT: BRCA C-terminal; HD: Hydrophobic domain; MIT: microtubule-interacting and trafficking domain; MBD: microtubule-binding domain; AAA: ATPase associated with diverse cellular activities; NBD: nucleotide-binding domain; HBD: helix-bundle domain. The G576 and K577 residues highlighted in red are essential for ATPase activity, and they were mutated to make dead mutant of TbSPA for *in vitro* microtubule-severing assay. (**B**). The AAA domain and the BRCT domain in TbSPA predicted by SWISS-MODEL. The templates used for modeling are 5Z6Q and 3AL2. (**C**). Purification of TbSPA truncation (a.a. 457-814) proteins for *in vitro* microtubule-severing assay. GA/AA stands for the G576A and K577A double mutations. (**D**). Effect of purified TbSPA truncation protein and its GK/AA mutant on microtubule length. MT: microtubule. (**E**). Effect of purified TbSPA truncation protein and its GK/AA mutant on microtubule integrity. Shown is the quantitation of the percentage of defective microtubules after incubation with purified TbSPA or its GK/AA mutant. Error bars indicate S.D. from three independent experiments. ns, no significance; ****, *p*<0.001. (**F**). Effect of purified TbSPA truncation protein and its GK/AA mutant on microtubule integrity. The yellow asterisks indicate the individual microtubules used for measurement of fluorescence signal intensity, which were presented below the images. Arrows and brackets indicate nicks (weak fluorescence signal) on the microtubules.

We performed *in vitro* microtubule severing assay using purified recombinant TbSPA-truncation protein containing the AAA domain and MBD domain and its activity-dead mutant bearing point mutations (G576A and K577A, abbreviated as GK/AA) at the Walker A motif of the nucleotide-binding domain (Fig. 1C). Rhodamine-labeled microtubules were assembled *in vitro*. Incubation of wild-type and mutant TbSPA-truncation proteins had no effect on the length of microtubules (Fig. 1D). The wild-type protein, but not the mutant protein, caused severe damages to the integrity of microtubules (Fig. 1E, F), in which microtubules did not show uniform fluorescence intensity along their length (Fig. 1F). It appears that the recombinant TbSPA-truncation protein did not severe microtubules into shorter fragments, but instead generated nicks on the microtubule lattice, which had lower fluorescence intensity (Fig. 1F, arrows and brackets). Nonetheless, these results suggest that TbSPA possesses microtubule severing activity *in vitro*.

### TbSPA localizes to the nucleus and associates with spindle during mitosis

We determined the subcellular localization of TbSPA during the entire cell cycle by immunofluorescence microscopy with cells expressing endogenously epitope-tagged TbSPA (Fig. 2A). During interphase (G1, S and G2 phases), TbSPA appeared to concentrate at the periphery of the nucleolus (Fig. 2A). When cells entered metaphase and anaphase A, TbSPA was concentrated around the mitotic spindle, which was labeled with anti-HA antibody for the endogenously 3HA-tagged β-tubulin (Fig. 2A). In anaphase B when the spindle was elongated, TbSPA was concentrated near the two spindle poles, and at telophase when spindle was disassembled, TbSPA was enriched at the periphery of the nucleolus (Fig. 2A).

**Figure 2.**
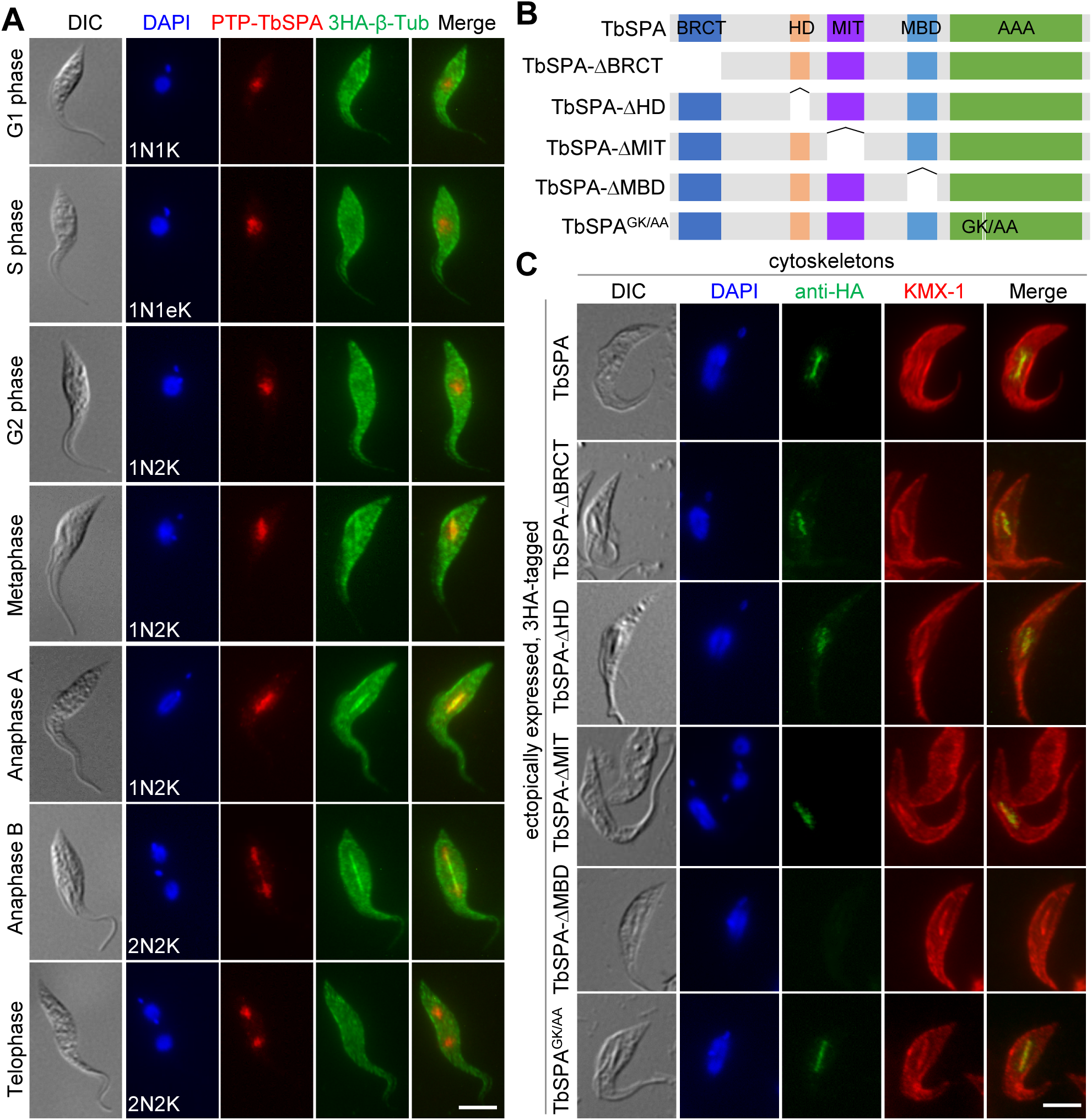
Subcellular localization of TbSPA and the structural motifs required for TbSPA association with spindle microtubules. (**A**). Immunofluorescence microscopic analysis of PTP-TbSPA and 3HA-β-Tub during different cell cycle stages in procyclic trypanosomes. Scale bar: 5 μm. (**B**). Schematic illustration of the structural motifs of TbSPA and the deletion mutants of TbSPA used for ectopic expression. (**C**). Immunofluorescence microscopic analysis of 3HA-tagged wild-type, domain-deletion mutants, and GK/AA mutant of TbSPA in detergent-extracted cytoskeletons. Scale bar: 5 μm.

We next examined the potential requirement of the structural domains and the microtubule-severing activity of TbSPA for subcellular localization. We ectopically overexpressed TbSPA, its domain-deletion mutants, and its activity-dead mutant or the GK/AA mutant (Fig. 2B) in trypanosomes, and examined their localization in intact cells and detergent-extracted cytoskeletons by immunofluorescence microscopy (Figs. S1A and 2C). To minimize the disruptive effect of overexpressed TbSPA on spindle integrity, cells were induced with tetracycline for 2 hours, at which there was no significant effect on spindle (see Fig. 5B below). In intact cells, the wild-type and mutant TbSPA proteins all localized around the spindle (Fig. S1A), similar to the endogenously PTP-tagged TbSPA (Fig. 2A). In detergent-extracted cytoskeletons, TbSPA, TbSPA-ΔBRCT, TbSPA-ΔHD, TbSPA-ΔMIT, and TbSPA^GK/AA^ were all detected around the spindle (Fig. 2C), but TbSPA-ΔMBD was undetectable (Fig. 2C). It suggests that TbSPA-ΔMBD was able to localize to spindle periphery but lost binding capability to spindle microtubules. Overall, these results demonstrated that TbSPA binds to spindle microtubules during mitosis and this microtubule-binding capability depends on the MBD domain.

### Overexpression of TbSPA inhibits cell proliferation and disrupts chromosome segregation

When TbSPA tagged with a C-terminal triple HA epitope was ectopically overexpressed under a tetracycline-inducible promoter in the procyclic form, we observed a strong growth defect, which was dependent on the dose of tetracycline (Fig. 3A) and the amount of overexpressed TbSPA-3HA protein (Fig. 3B). Overexpression of TbSPA without any epitope tag also caused severe growth defects (Fig. 3C). These results suggest a dominant effect caused by TbSPA overexpression. We further asked whether overexpression of the domain-deletion mutants and the activity-dead mutant of TbSPA would affect the dominant effect of TbSPA. Ectopic overexpression of the BRCT domain-deletion mutant, TbSPA-ΔBRCT, caused severe growth defects, and overexpression of the HD-deletion mutant, TbSPA-ΔHD, caused moderate growth defects (Fig. 3D), whereas overexpression of the MIT domain-deletion mutant, TbSPA-ΔMIT, the MBD domain-deletion mutant, TbSPA-ΔMDB, or the GK/AA mutant exerted little effects on cell growth (Fig. 3D). These results suggest that the MIT domain, the MBD domain, and the microtubule-severing activity of TbSPA are each required for TbSPA function.

**Figure 3.**
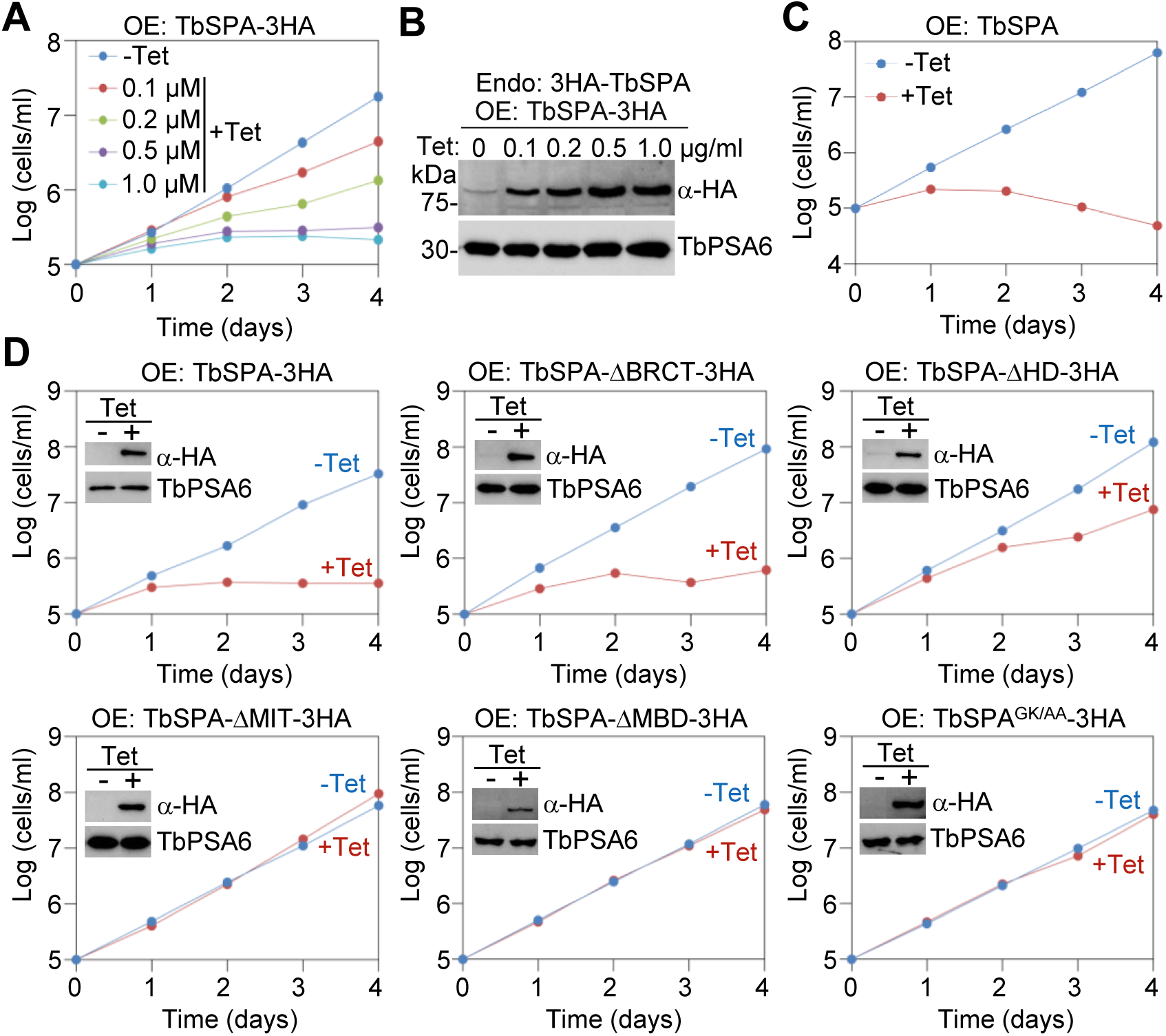
Overexpression of TbSPA in the procylic form of *T. brucei* inhibits cell proliferation. (**A**). Growth curves of procyclic trypanosomes overexpressing TbSPA-3HA induced with different concentrations of tetracycline. (**B**). Western blots of endogenous 3HA-TbSPA and ectopically overexpressed TbSPA-3HA induced with different concentrations of tetracycline. TbPSA6, the 20S proteasomal α-6 subunit, served as the loading control. (**C**). Growth curve of procyclic trypanosomes overexpressing TbSPA without an epitope tag. (**D**). Growth curves of procyclic trypanosomes overexpressing 3HA-tagged TbSPA, its various domain-deletion mutants, or the GK/AA mutant. Insets show the western blots of TbSPA-3HA and its mutants induced with 1.0 µg/ml tetracycline and western blots of TbPSA6 serving as the loading control.

Since ectopically overexpressed TbSPA localized to the spindle (Fig. 2C), we investigated the potential effects of TbSPA overexpression on mitosis. Quantitation of cells with different numbers of nuclei (N) and kinetoplasts (K) showed that after induction of TbSPA overexpression for 8 h, there was a significant reduction of 1N1K cells and 2N2K cells and a significant increase of 1N2K cells, and anucleate (0N1K) cells (Fig. 4A). After TbSPA overexpression for 16 h, there was a significant increase of uni-nucleated cells with either a large or a small nucleus, which were named 1N*1K cells (Fig. 4A). There was no accumulation of xNxK (x>2) cells after induction of TbSPA overexpression after up to 24 h (Fig. 4A). Thus, TbSPA overexpression did not impair cytokinesis, but disrupted mitosis. The accumulation of uni-nucleated cells with an abnormal-sized nucleus was further confirmed by measurement of the nucleus surface size, which showed a wider distribution of nucleus size upon TbSPA overexpression (Fig. 4B). In control uni-nucleated cells, the nucleus size was in the range of 4 to 8 μm^2^, whereas upon TbSPA overexpression for 16 h, cells with the nucleus size smaller than 4 μm^2^ or larger than 8 μm^2^ were significantly increased, following a corresponding decrease of cells with the nucleus size between 4-6 μm^2^ (Fig. 4C). Those cells with an abnormal-sized nucleus (Fig. 4D) apparently were derived from the 2N2K cells that have undergone an asymmetrical nuclear division, which generated a small nucleus and a large nucleus (Fig. 4D). The asymmetrical nuclear division was further confirmed by FISH using a megabase chromosome probe and a mini-chromosome probe, which showed that asymmetrical nuclear division generated two nuclei of different size (Fig. 4E, F).

**Figure 4.**
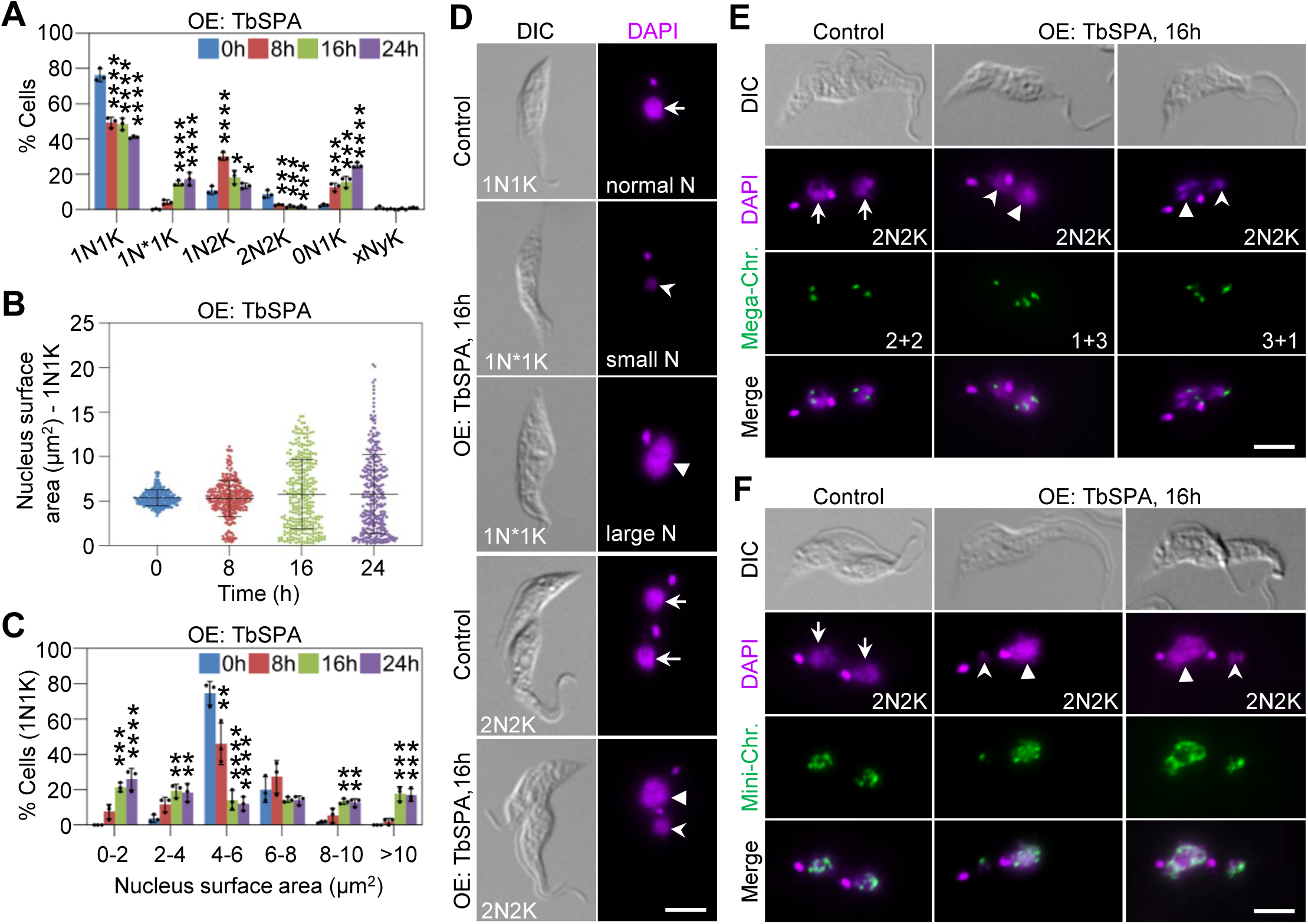
Overexpression of TbSPA disrupts chromosome segregation. (**A**). Effect of TbSPA overexpression on cell cycle progression. Shown is the quantitation of cells with different numbers of nuclei (N) and kinetoplasts (K) before and after TbSPA overexpression. Error bars indicate S.D. from three independent experiments. *, *p*<0.05; ***, *p*<0.001; ****, *p*<0.0001. (**B**). Measurement of nucleus size of 1N1K cells before and after TbSPA overexpression. (**C**). Quantitation of cells with different nucleus sizes before and after TbSPA overexpression. Error bars indicated S.D. from three independent experiments. **, *p*<0.01; ***, *p*<0.001; ****, *p*<0.0001. (**D**). Effect of TbSPA overexpression on nucleus size. Arrows: normal-sized nucleus; Open arrowheads: small nucleus; Solid arrowheads: large nucleus. Scale bar: 5 μm. (**E**). Segregation of mega-chromosomes detected by FISH in control and TbSPA overexpression cells. Arrows: normal-sized nucleus; Open arrowheads: small nucleus; Solid arrowheads: large nucleus. The numbers indicate the distribution of the FISH-detected chromosomes between the two segregated nuclei. Scale bar: 5 μm. (**F**). Segregation of mini-chromosomes detected by FISH in control and TbSPA overexpression cells. Arrows: normal-sized nucleus; Open arrowheads: small nucleus; Solid arrowheads: large nucleus. Scale bar: 5 μm.

### Overexpression of TbSPA impairs spindle integrity

Because TbSPA is a microtubule-severing enzyme (Fig. 1E, F) and associates with mitotic spindle (Fig. 2), we wondered whether its overexpression disrupted spindle integrity, thereby causing asymmetrical nuclear division (Fig. 4). Using endogenously 3HA-tagged β-tubulin and the anti-β-tubulin antibody KMX-1 as spindle markers, we examined the mitotic spindle in cells overexpressing TbSPA. We found that the 1N2K cells with a detectable spindle were significantly reduced by ∼36% and ∼59% after TbSPA overexpression for 4 h and 6 h, respectively (Fig. 5A, B). Notably, in these spindle-positive cells, ∼56% and ∼82% of them contained a damaged spindle with weaker fluorescence signal (Fig. 5A, B), suggestive of reduced numbers of microtubules and an increase of broken (severed) microtubules.

**Figure 5.**
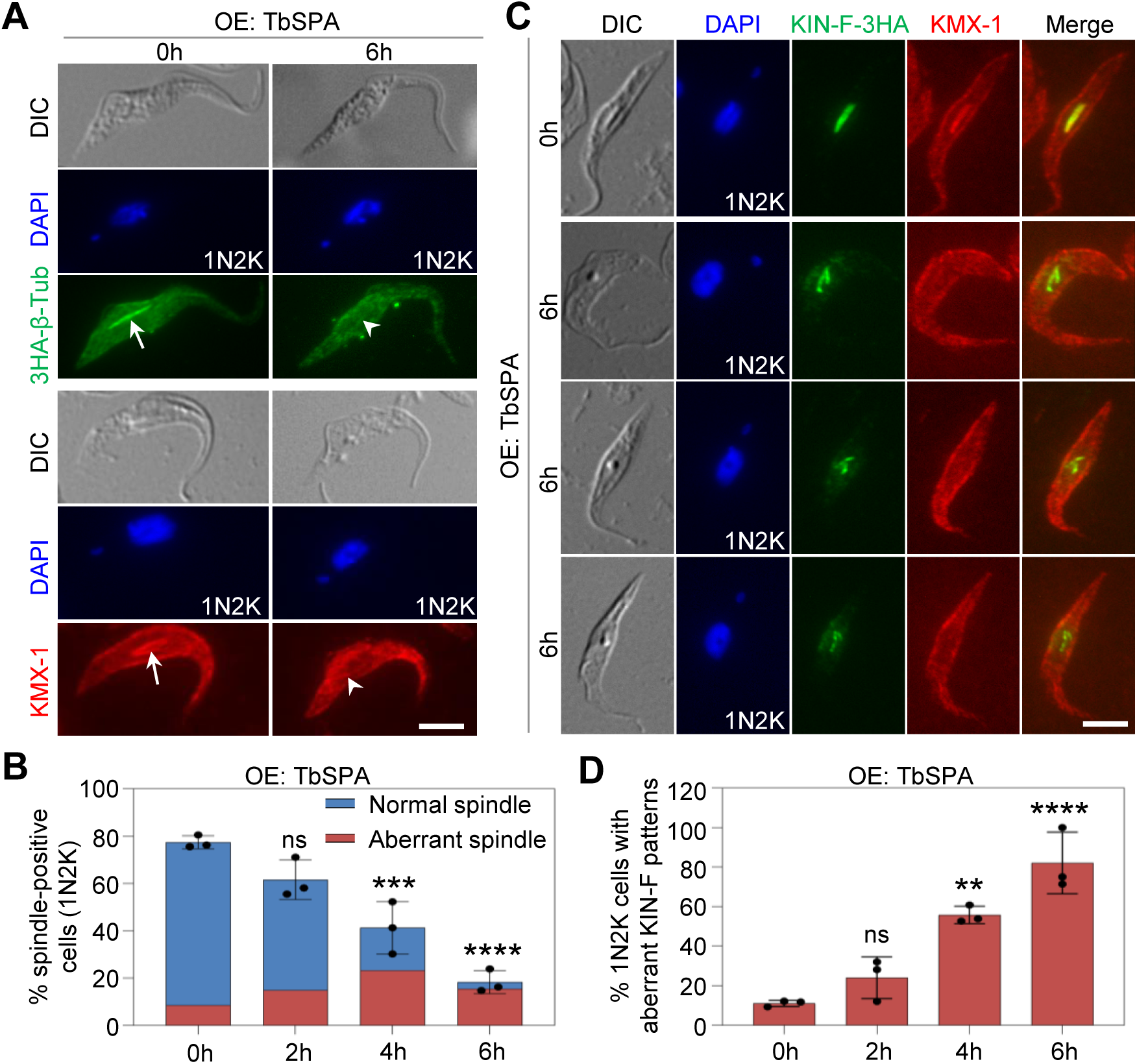
Overexpression of TbSPA disrupts spindle integrity. (**A**). Immunofluorescence microscopic analysis of spindle in control and TbSPA overexpression cells. Spindle was detected by either anti-HA antibody to label endogenously 3HA-tagged β-tubulin (β-Tub) or the KMX-1 antibody to label native β-tubulin. Arrows: normal spindle; Open arrowheads: defective spindle. Scale bar: 5 μm. (**B**). Quantitation of cells with spindle and defective spindle before and after TbSPA overexpression. Error bars indicate S.D. from three independent experiments. (**C**). Effect of TbSPA overexpression on the localization of KIN-F, a spindle-associated orphan kinesin. Scale bar: 5 μm. (**D**). Quantitation of cells with aberrant localization patterns of KIN-F before and after TbSPA overexpression. Error bars indicate S.D. from three independent experiments.

To further corroborate this observation, we used the spindle-associated kinesin protein KIN-F as a marker to examine the spindle integrity. In the detergent-extracted cytoskeletons of control cells, KIN-F associated with the entire spindle (Fig. 5C), whereas in the detergent-extracted cytoskeletons of TbSPA-overexpressed cells, KIN-F displayed various abnormal-shaped patterns (Fig. 5C), and in these cells the spindle had weak signal or were detected as discontinued punctate dots by the KXM-1 antibody (Fig. 5C). The 1N2K cells with aberrant KIN-F signal patterns were significantly increased by ∼45% and ∼71% after TbSPA overexpression for 4 h and 6 h, respectively (Fig. 5D). Altogether, these results demonstrated that TbSPA overexpression disrupted spindle integrity by either eliminated the spindle or damaged some of the spindle microtubules.

### Knockdown of TbSPA in the procyclic form of *T. brucei* impairs spindle integrity

We performed RNAi to characterize TbSPA function in the procyclic form using the Stem-loop RNAi construct. Western blotting showed the gradual knockdown of the endogenously PTP-tagged TbSPA protein to undetectable levels after RNAi induction for 7 days (Fig. 6A). This knockdown caused weak growth defects after 4 days of RNAi induction, but it did not inhibit cell proliferation (Fig. 6B). We further attempted to knock out both alleles of the *TbSPA* gene, and we were able to knockout one allele, but were not able to obtain viable cells when attempting to delete the second allele (Fig. S1B), suggesting that *TbSPA* is essential for cell viability. Therefore, we used the TbSPA RNAi cell line to characterize the potential effect of TbSPA depletion on cell cycle progression in the procyclic form. Given that TbSPA associates with spindle, we examined the morphology and/or integrity of spindle by immunofluorescence microscopy with the KMX-1 antibody, which showed that in metaphase and anaphase-A cells, but not anaphase-B cells, the spindle appeared to have missing fluorescence signal at certain locations (Fig. 6C), suggestive of disrupted integrity of spindle microtubules during early stages of mitosis. After RNAi induction for 4 days, the metaphase and anaphase-A cells with spindle defects were significant increased by ∼26% and ∼9%, respectively (Fig. 6D). To further characterize the spindle defects, we used KIN-F as another spindle marker in immunofluorescence microscopy, which also detected abnormal patterns of KIN-F localization on the spindle (Fig. 6E). Such cells with aberrant KIN-F patterns increased by ∼20% in metaphase cells and by ∼29% in anaphase-A cells (Fig. 6F). These results suggest that knockdown of TbSPA caused spindle defects, likely by disrupting microtubule dynamics.

**Figure 6.**
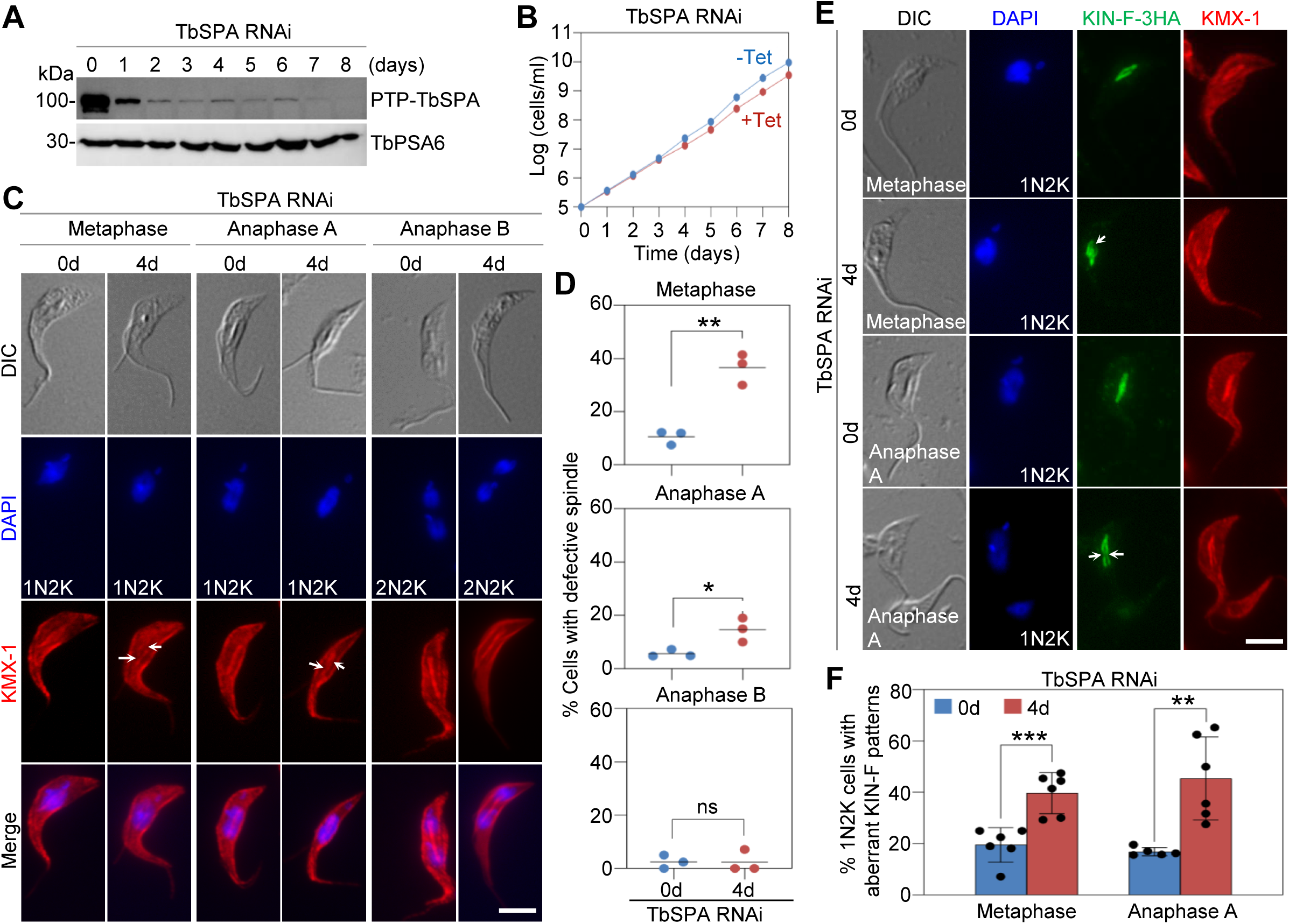
Knockdown of TbSPA in the procyclic form of *T. brucei* impairs spindle structural integrity. (**A**). Western blotting to detect the levels of TbSPA after TbSPA RNAi induction in the procyclic form. TbPSA6 served as a loading control. (**B**). Growth curves of non-induced and RNAi induced TbSPA RNAi cell line of the procyclic form. (**C**). Effect of TbSPA knockdown on spindle structural integrity. Arrows indicate missing parts of the spindle microtubules. Scale bar: 5 μm. (**D**). Quantitation of mitotic cells with defective spindle structure. Error bars indicate S.D. from three independent experiments. (**E**). Effect of TbSPA knockdown on the localization of the spindle-associated orphan kinesin KIN-F. Cells were co-immunostained with the KMX-1 antibody for spindle and the anti-HA antibody for KIN-F-3HA. Arrows indicate missing parts of the KIN-F signal on the spindle. Scale bar: 5 μm. (**F**). Quantitation of 1N2K cells with aberrant KIN-F patterns in control and TbSPA RNAi-induced cells. Error bars indicate S.D. from six independent experiments. **, *p*<0.01; ***, *p*<0.001.

### Knockdown of TbSPA causes defective chromosome segregation

We investigated the effect of TbSPA knockdown on chromosome segregation by monitoring the segregation of kinetochores using the kinetochore protein KKT2 as a marker in immunofluorescence microscopy. In control cells at metaphase, the kinetochores were aligned at the metaphase plate, and frequently only two parallel foci were detected on the two lines of spindle microtubules (Fig. 7A). In TbSPA RNAi cells during metaphase, however, the kinetochores appeared to be misaligned, with more than two foci detected along the somewhat defective spindle microtubules (Fig. 7A). The metaphase cells with mis-aligned kinetochores were increased by ∼33% after RNAi induction for 4 days (Fig. 7B). In control cells during anaphase A, kinetochores segregated to the two spindle poles (Fig. 7A), whereas in TbSPA RNAi cells during anaphase A, some kinetochores did not segregate to the poles but were lagging behind (Fig. 7A). The anaphase-A cells with lagging kinetochores were increased by ∼34% after RNAi induction for 4 days (Fig. 7B). However, there was no significant increase of anaphase-B cells with lagging kinetochores (Fig. 7B), in agreement with the insignificant change of the anaphase-B cells with defective spindle (Fig. 6D). These results demonstrated the involvement of TbSPA in promoting chromosome alignment in metaphase and chromosome segregation in anaphase A.

**Figure 7.**
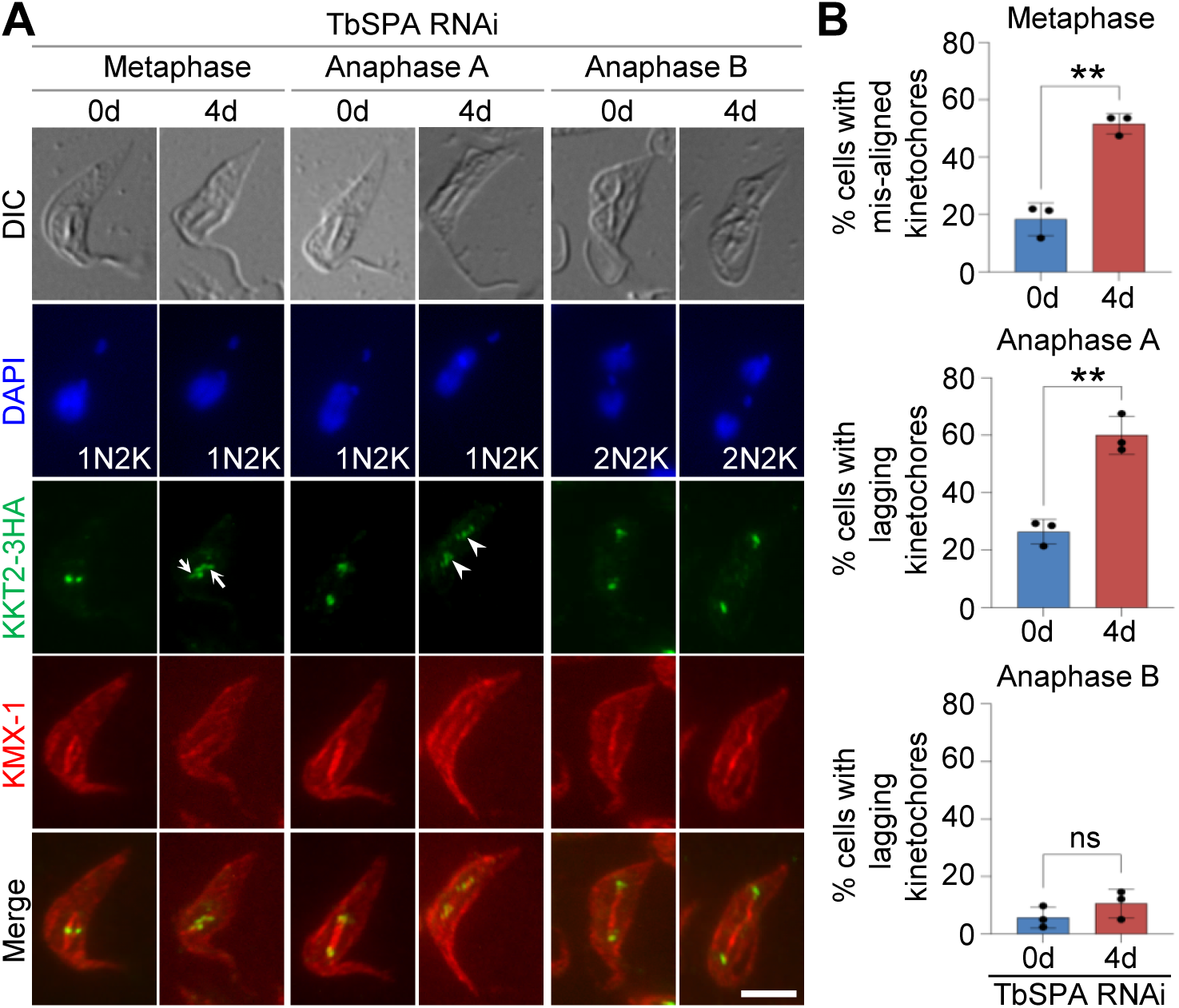
Knockdown of TbSPA disrupts chromosome segregation. (**A**). Effect of TbSPA RNAi on the segregation of chromosomes. Shown are the immunostainings of kinetochore protein KKT2 in control and TbSPA RNAi-induced cells. Arrows: mis-aligned kinetochores; Open arrowheads: lagging kinetochores. Scale bar: 5 μm. (**B**). Quantitation of metaphase cells with mis-aligned kinetochores and anaphase cells with lagging kinetochores in control and TbSPA RNAi-induced cells. Error bars indicate S.D. from three independent experiments. **, *p*<0.01; ns, no significance.

We further characterized the mitotic defects of TbSPA RNAi cells by time-lapse video microscopy of live cells using mCherry-tagged histone H2B to label the chromosomes (Fig. 8A and Videos S1-S3). Control and TbSPA RNAi cells were imaged under a confocal microscope for up to 15 h, and cells that were able to complete mitosis or that had started, but were not able to complete mitosis within this time frame were all included in the analysis. Figure 8A shows a control cell that completed mitosis, a TbSPA RNAi cell that failed mitosis, and a TbSPA RNAi cell that underwent slower mitosis. With the time of metaphase (before anaphase onset) set at minute 0, the control cell entered anaphase A at minute 6 and anaphase B at minute 12, and initiated cytokinesis at minute 64 (Fig. 8A and Video S1). The TbSPA RNAi cell that failed mitosis entered anaphase A at minute 6, but did not progress into anaphase B and initiated cytokinesis at minute 84 (Fig. 8A and Video S2). The TbSPA RNAi cell that had slower mitosis entered anaphase A at minute 6 and anaphase B at minute 16, and initiated cytokinesis at minute 94 (Fig. 8A and Video S3). The TbSPA RNAi cells had no problems entering anaphase A, but they either were not able to progress into anaphase B or took longer time to enter anaphase B. Further, mitotic progression in control and TbSPA RNAi cells was quantitatively analyzed by measuring the nucleus distance from 60 min prior to anaphase onset (or the end of metaphase) and until 80 min after anaphase onset (Fig. 8B). In the control cell, the nucleus distance doubled after ∼28 min from anaphase onset and remained until cytokinesis (Fig. 8A, B). In the TbSPA RNAi cell that failed mitosis, the nucleus distance was first increased by ∼37% after ∼6 min from anaphase onset, but then started to reduce after 22 min and further reduced by ∼24% after 52 min until cytokinesis (Fig. 8B). Those cells that failed mitosis increased significantly from ∼3% to ∼20% after TbSPA RNAi for 4 days (Fig. 8C). In the TbSPA cell that had slower mitosis, the nucleus distance had similar pattern to that of the control cell (Fig. 8B), despite delays in anaphase and cytokinesis initiation (Fig. 8A). In these TbSPA RNAi cells that had slower mitosis, the average mitosis time was ∼99 min, which was 53% longer than the average mitosis time (∼65 min) of the control cells (Fig. 8D). These results suggest that TbSPA knockdown impaired chromosome segregation, causing delays in anaphase onset and/or progression, which further delayed the completion of mitosis and the initiation of cytokinesis.

**Figure 8.**
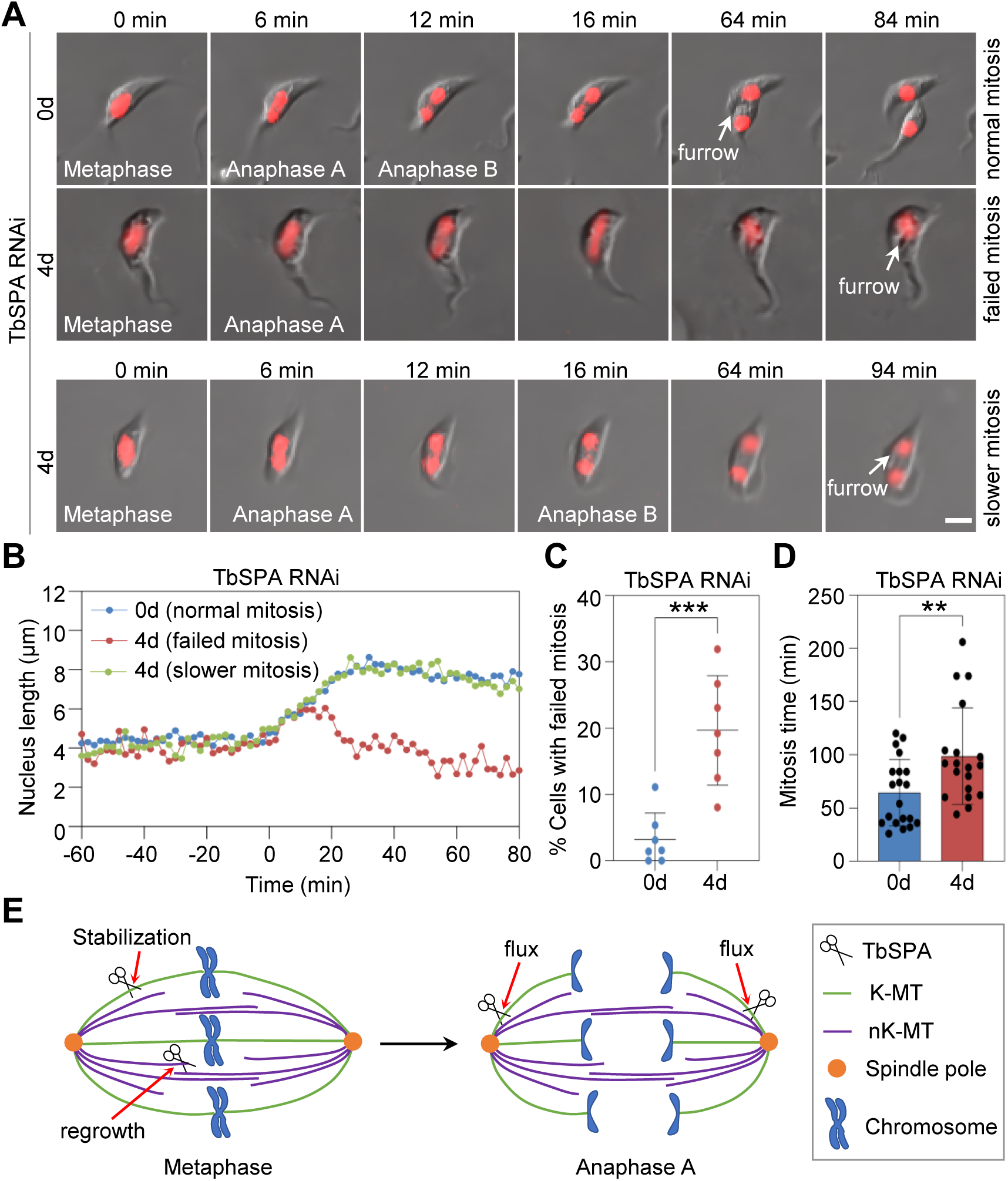
Knockdown of TbSPA disrupts nuclear division. (**A**). Selected images from time-lapse videos of control and TbSPA RNAi-induced cells. Nucleus was labeled with mCherry-tagged histone H2B. Scale bar: 5 μm. (**B**). Measurement of nucleus length of control and TbSPA RNAi-induced cells from the images of the time-lapse videos. (**C**). Quantitation of cells with failed mitosis in control and TbSPA RNAi-induced cells. Shown are the quantitation of live-imaged cells that initiated but failed mitosis, as shown in panel A. Error bars indicate S.D. from six independent experiments. ***, *p*<0.001. (**D**). Quantitation of mitosis time in control and TbSPA RNAi-induced cells. Error bars indicate S.D. **, *p*<0.01. (**E**). Model to explain the role of TbSPA in regulating spindle dynamics and chromosome movement.

## Discussion

Microtubule-severing enzymes, including katanin, spastin, and fidgetin, regulate diverse cellular processes, including mitosis and cytokinesis, in eukaryotes. Unlike microtubule depolymerases that depolymerize microtubules from their ends (29), microtubule-severing enzymes generate internal breaks in the microtubule lattice (30). These enzymes are characterized by the presence of a C-terminal AAA-type ATPase domain that catalyzes microtubule severing to regulate microtubule dynamics. The parasitic protozoan *T. brucei* has all the three types of microtubule-severing enzymes, but none of them have been found to play roles in mitosis in previous studies. We have revisited the localization and function of the *T. brucei* spastin homolog TbSPA in the procyclic form, and we showed that TbSPA localizes to the nucleus, associates with the mitotic spindle, and regulates spindle dynamics to promote chromosome segregation in the insect life cycle stage. In the mammalian bloodstream stage of the *T. brucei* life cycle, however, TbSPA appears to promote cell abscission during cytokinesis, likely by severing the subpellicular microtubules located at the cytoplasmic bridge connecting the two daughter cells, without detectable roles in mitosis (22). Therefore, TbSPA plays life cycle form-specific functions by promoting mitosis in the procyclic form and cytokinesis in the bloodstream form.

We determined the structural requirement for TbSPA localization and function in the procyclic form of *T. brucei* and identified features distinct from its human counterpart. The N-terminal BRCT domain (Fig. 1A, B), which is not present in the spastin homologs from other organisms (28), is not required for TbSPA localization and function (Figs. 2 and 3). The MBD domain, as expected, is required for the association of TbSPA with spindle microtubules and is essential for TbSPA function, although it is not required for TbSPA localization to the nucleus (Figs. 2, 3 and S1A). The MIT domain is essential for TbSPA function, but not for TbSPA localization (Figs. 2 and 3), in contrast to the MIT domain in human spastin that targets the protein to the midbody and endosomes through interaction with the ESCRT-III proteins IST1 or CHMP1B (28). The HD domain in TbSPA is important, but not essential, for function, and is not required for TbSPA localization (Figs. 2 and 3), which also differs from the function of the HD domain in human spastin that targets the protein to the ER due to its affinity to the membrane (28). TbSPA differs substantially from its human counterpart in the regulation of its subcellular localization through the conserved structural domains.

We also demonstrated that the recombinant TbSPA truncation protein containing the AAA domain and the MBD domain generates nicks on the microtubule lattice *in vitro* (Fig. 1E, F), but it does not cut microtubules into shorter fragments (Fig. 1D). This could be attributed to the weaker enzymatic activity of the purified recombinant protein than the native protein in trypanosome cells or due to the weaker enzymatic activity in our *in vitro* assays. Previous studies showed that human spastin generates nicks by extracting tubulins from the microtubule lattice, and then new tubulin proteins are incorporated into the nicks, thereby stabilizing the microtubules (31,32). Thus, recombinant TbSPA appears to possess activities comparable to the recombinant human spastin. Mutation of the Walker A motif in the AAA domain disrupted the activity of TbSPA (Fig. 1E, F), suggesting that this *in vitro* activity is ATP-dependent. These results demonstrated TbSPA is a conserved microtubule-severing enzyme that regulates microtubule dynamics.

Overexpression of TbSPA and knockdown of TbSPA in the procyclic form both disrupt spindle microtubule architecture, with the former destroying the spindle and the latter generating a defective spindle with loss of microtubule integrity and abnormal morphology (Figs. 5 and 6). Given TbSPA’s enzymatic activity in severing microtubules (Fig. 1E, F), one would expect that knockdown of TbSPA should stabilize spindle microtubules. This paradoxical observation reflects the mechanistic function of microtubule-severing enzymes, including spastin and katanin, in regulating microtubule dynamics, and similar observations have been made in previous studies, which reported that deficiency in these enzymes leads to reduced microtubule mass and causes loss and defects of microtubules (33–37). Spastin is not playing a sole, destructive role in disassembling microtubules, but instead functions as a dual-function enzyme by severing microtubules and promoting microtubule regrowth and stability (31,32). This dual function of spastin increases microtubule number and mass, thereby regulating microtubule architecture and dynamics (31,32). In light of this role of spastin in regulating microtubule dynamics, we speculate that TbSPA plays a similar role in severing spindle microtubules and promoting microtubule regrowth and stability to establish a bipolar spindle (Fig. 8E). In the absence of TbSPA, some spindle microtubules disassemble due to instability, and some other microtubules fail to elongate due to inability of regrowth, thus producing a defective spindle (Fig. 6C, D). Dynamic instability occurred on kinetochore microtubules can cause kinetochore-microtubule detachment, leading to mis-aligned kinetochores in metaphase and lagging kinetochores in anaphase A, as observed in TbSPA RNAi cells (Fig. 7). Notably, no lagging kinetochores were detected in anaphase-B cells after TbSPA RNAi (Fig. 7), suggesting that the RNAi cells were able to correct the kinetochore-microtubule attachment errors. These attachment errors can activate mitotic checkpoints to delay mitosis, allowing more time for the cells to correct the errors. The significant delays in mitosis in some TbSPA RNAi cells observed by time-lapse video microscopy (Fig. 8 and Videos S1-S3) suggest potential checkpoint activation by TbSPA deficiency-induced spindle microtubule-kinetochore attachment errors in *T. brucei*.

The mechanistic role of spastin in promoting chromosome movement during anaphase in animal cells is through the severing of spindle microtubules from the minus ends, generating the so-called poleward flux to move chromosomes towards the spindle poles in anaphase A (19). This process is facilitated by microtubule depolymerization catalyzed by kinesin-13 from the minus ends, which shortens the kinetochore microtubules (38). Because of the observed chromosome segregation defects during anaphase A in TbSPA RNAi cells (Figs. 7 and 8), we postulate that TbSPA also plays a role in promoting chromosome segregation through the poleward flux mechanism, likely by severing spindle microtubules from their minus ends at the spindle poles (Fig. 8E). This can also be facilitated by KIN13-1/Kif13-1, a kinesin-13 family protein localizing to the mitotic spindle and playing essential roles in chromosome segregation (20,21). In this regard, *T. brucei* appears to employ a conserved mechanism for chromosome movement during anaphase A through the cooperative action of spastin and kinesin-13. Future work will be directed to explore how TbSPA, Kif13-1/KIN13-1, and other spindle-associated proteins may cooperate to regulate spindle dynamics in *T. brucei*, which will shed lights on the regulation of mitosis in the absence of the evolutionarily conserved spindle motor and kinetochore motor proteins.

## Materials and Methods

### Trypanosome cell culture

Three strains of the procyclic (insect) form of *T. brucei*, the 29-13 strain (39), the SmOx strain (40) and the Lister427 strain, were used in this study. The 29-13 strain was cultured in SDM-19 medium containing 10% heat-inactivated fetal bovine serum (Millipore-Sigma), 15 µg/ml G418, and 50 µg/ml hygromycin at 27°C, the SMOX strain was cultured in SDM-79 medium containing 10% heat-inactivated fetal bovine serum (Millipore-Sigma) and 50 µg/ml hygromycin at 27°C, and the Lister427 strain was cultured in SDM-79 medium containing 10% fetal bovine serum at 27°C. Cells were routinely diluted with fresh medium every 3 days or whenever the cell density reaches 5 × 10^6^ cells/ml.

### Ectopic overexpression, RNA interference, and knock-out of *TbSPA* gene

To ectopically overexpress TbSPA and its mutants in *T. brucei*, the full-length coding sequence of TbSPA gene was PCR amplified from the genomic DNA and cloned into the pLew100-BLE vector (39). DNA fragments encoding each of the four domain-deletion mutants of TbSPA were obtained by PCR and similarly cloned into the pLew100-BLE vector. To make the G576A and K577A double mutation of TbSPA, site-directed mutagenesis was performed using the pLew100-TbSPA-BLE plasmid as the template, and mutation of the two sites were confirmed by sequencing. Each of these plasmids was linearized by restriction digestion with NotI and used to transfect the 29-13 strain by electroporation. Successful transfectants were selected with 2.5 µg/ml phleomycin and cloned by limiting dilution in a 96-well plate with the SDM-79 medium containing 20% heat-inactivated fetal bovine serum, 2.5 µg/ml phleomycin, 15 µg/ml G418, and 50 µg/ml puromycin. Overexpression of TbPSA and its mutants was induced with 1.0 µg/ml tetracycline.

To generate TbSPA RNAi cell line, a stem-loop-based RNAi construct was generated by cloning a 700-bp DNA fragment (nucleotides 777-1,476) of the *TbSPA* gene into the HindIII/XhoI sites of the pSL vector (41), and subsequently the same DNA fragment was cloned in an opposite direction into the AflII/BamHI sites of the same pSL vector. The resulting plasmid, pSL-TbSPA, was linearized by NotI restriction digestion and then used for transfection into the SmOx by electroporation. Transfectants were selected with 2.5 µg/ml phleomycin and cloned by limiting dilution in a 96-well plate containing the SDM-79 medium, 20% heat-inactivated fetal bovine serum, 2.5 µg/ml phleomycin, and 50 µg/ml puromycin. RNAi was induced by incubating the cells with 1.0 µg/ml tetracycline.

Knockout of each of the two alleles of the TbSPA gene was carried out by homologous recombination-based gene replacement. 100-bp fragment of the 5’UTR and 100-bp fragment of the 3’UTR of TbSPA gene were included in the PCR primers to amplify the puromycinN-acetyltransferase (pac) gene. The PCR products were purified, and used to transfect the *T. brucei* 29-13 strain by electroporation. Transfectants were selected with 1 μg/ml puromycin and then cloned by limiting dilution, as described above. Genomic DNA from the clonal cell lines were extracted and used to confirm the replacement of one TbSPA allele by PCR. The *TbSPA*^+/-^ cell lines were then used to replace the second TbSPA allele using the same approach, by replacing it with the blasticidin-S deaminase gene. Transfectants were selected with 10 μg/ml blasticidin. No viable cells were obtained, despite numerous attempts, suggesting that knockout of *TbSPA* is lethal in the procyclic form.

### *In situ* epitope tagging of proteins

One of the respective endogenous loci was tagged by either PCR-based epitope tagging method (42) or plasmid-based method using the tagging plasmids developed in the Gunzl lab (43). To procced with PCR-based epitope tagging, PCR products were purified from agarose gel and 4 μg of DNA was used for transfection by electroporation. For plasmid-based method, 15 μg of plasmid DNA was linearized with appropriate restriction enzymes, precipitated, and used for electroporation. Transfectants were selected with 10 μg/ml blasticidin, 15 μg/ml G418, or 1 μg/ml puromycin, accordingly, and were further cloned by limiting dilution as described above.

### Expression and purification of recombinant spastin

The C-terminal fragment of *TbSPA* gene (nucleotides 1,339-2,445), which encodes the MBD and the AAA domains, was PCR amplified and cloned into pGEX-4T-3 to express GST-TbSPA(457-814). The resultant vector was sequenced and used as a template for site-directed mutagenesis to mutate G576A/K577A, generating a catalytically inactive mutant GST-TbSPA(457-814)^GK/AA^. The plasmids were each transformed into *E. coli* strain BL-21, and cells were induced to express the recombinant protein with 0.03 mM IPTG for 16 h at 16°C. Cells expressing the recombinant protein were resuspended in 0.1% Triton X-100 in PBS, lysed by sonication, and the lysate was cleared by centrifugation at 20,627 *g* for 10 min at 4°C. Cleared lysate was incubated with the Glutathione Sepharose 4B beads, pre-equilibrated with the same buffer, for 1 h at 4°C with gentle agitation. Beads were washed with 0.1% Triton X-100 in PBS, and bound proteins were eluted with 20 mM glutathione in 50 mM Tris, pH 9.0, 0.1% Triton X-100, 100 mM NaCl, and 1 mM DTT. Eluted proteins were concentrated and buffer-exchanged with Amicon Ultra Centrifugal Filters 10K (Millipore-Sigma).

### Tubulin polymerization and microtubule severing assay

*In vitro* tubulin polymerization using porcine brain tubulin was carried out as previously described (44). Briefly, non-labeled and rhodamine-labeled porcine brain tubulin (Cytoskeleton, Inc., Cat#: T240-B and TL590M-A, respectively) were mixed in BRB80-DTT buffer (80 mM K-PIPES, pH 6.8, 1.0 mM MgCl_2_, 1.0 mM EGTA, 1.0 mM DTT) supplemented with 1.0 mM Guanylyl-(α,β)-methylene-diphosphonate (GMP-CPP), and incubated for 5 min at 4°C, followed by ultracentrifugation at 353,000 *g* for 5 min at 4°C. The supernatant was transferred to a new tube, 1:4 diluted with BRB80-DTT, and incubated for 30 min at 37°C, to produce the microtubule seeds, which were pelleted at 353,000 *g* for 5 min at 27°C. The pellet was resuspended with BRB80-DTT and stored at room temperature. The microtubule seeds were then mixed with 20 μg of rhodamine-labeled porcine brain tubulin in the presence of 2.0 mM GTP and increased concentrations of Taxol, up to 10 μM, at 37°C. Microtubule polymerization was verified under the microscope and stored at room temperature.

Microtubule-severing assay was conducted in 0.2 ml tubes at room temperature. A 1:1 ratio (500 nM each) of purified recombinant TbSPA(457-814) and microtubules were mixed in BRB80-DTT, supplemented with Taxol and GTP, and in the presence of 1 mM ATP, for 15 min. The reaction was stopped by mixing 3 μl of the reaction solution with 1.8 μl of 4% PFA. Microtubules were immediately imaged with an inverted fluorescence microscope (Olympus IX71) with a PlanApo N 60× oil lens. Images were acquired using the Slidebook software (version 5.0; Intelligent Imaging Innovations) and analyzed with ImageJ.

### Immunostaining and immunofluorescence microscopy

*T. brucei* cells were washed once with PBS, resuspended in PBS, and attached to glass coverslips for 30 min at room temperature. For cytoskeleton preparation, the attached cells were treated with 1% Nonidet-P40 in PEME buffer (100 mM PIPES, pH 6.9, 2 mM EGTA, 1 mM MgSO_4_, and 0.1 mM EDTA) for 1 second. Cells and cytoskeletons were fixed with cold methanol for 10 min at -20°C, rehydrated with PBS, and incubated with blocking buffer (3% BSA in PBS) for 30 min at room temperature, followed by primary antibody incubation for 1 h at room temperature in the same buffer. The following primary antibodies were used: anti-HA monoclonal antibody (1:400 dilution), anti-Protein A polyclonal antibody (1:400 dilution), and anti-β-tubulin monoclonal antibody (KMX-1, 1:400 dilution). After 3 washes with PBS (5 min/wash under agitation), cells on the coverslip were incubated with FITC-conjugated or Alexa Fluor488-cojugated anti-mouse or anti-rabbit IgG (1:400 dilution), Alexa Fluor555-congugated anti-mouse IgG (1:400 dilution), or Cy3-conjugated anti-rabbit IgG (1:400 dilution). Cells were washed with PBS, mounted in VectaShield mounting medium, and stored at 4°C, protected from light, or imaged with an inverted fluorescence microscope (Olympus IX71) equipped with a cooled CCD camera and a PlanApo N 60x1.42 NA oil lens. For image acquisition, the Slidebook software was used, and images were processed using ImageJ.

### Fluorescence in situ hybridization (FISH)

The chromosome segregation was detected by FISH, as described (25). Briefly, the M5 ribosomal RNA gene on chromosome 8, which was amplified by PCR with the forward primer 5’-GGGTACGACCATACTTGGC-3’ and reverse primer 5’-AGAGTACAACACCCCGGGTT-3’, was used to label mega-chromosomes, and the mini-chromosomal 177-bpDNA repeat, which was amplified by PCR with the forward primer 5’-TAAATGGTTCTTATACGAATG-3’ and the reverse primer 5’AACACTAAAGAACAGCGTTG-3’, was used to label mini-chromosomes. Digoxigenin-dUTP (Sigma-Aldrich) was incorporated into the two DNA sequences during PCR. Purified PCR products were precipitated and resuspended in hybridization buffer (50% formamide, 2×saline-sodium citrate (SSC), 10% dextran sulfate).

Trypanosome cells attached to glass coverslips were fixed with cold methanol, rehydrated with PBS, prehybridized in the hybridization buffer for 60 min, and then denatured at 95°C for 5 min. Cells were probed with digoxigenin-labeled probes at 37°C for 16 h and washed extensively with wash buffer (three times with 2×SSC containing 50% formamide at 37°C, three times with 0.2% SSC at 50°C, three times with 4×SSC at room temperature). After blocking with 4×SSC containing 3% BSA for 30 min, cells were incubated with FITC-conjugated anti-digoxigenin antibody (1:400 dilution, Clone# DI-22, Catalog# F3523, Millipore-Sigma). After three washes with 2×SSC, cells were mounted in the mounting medium, and imaged under a fluorescence microscope. Images were acquired by the Slidebook software and processed with ImageJ.

### Time-lapse video microscopy

Histone H2B (Tb927.10.10460) was tagged with a C-terminal mCherry in TbSPA RNAi cell line by PCR-based epitope tagging method (42), and clonal cell lines obtained were used for time-lapse video microscopy. Non-induced and RNAi-induced (4 days) cells were attached to glass coverslips at room temperature. In the meantime, on a glass slide, a small chamber was made by four pieces of double-sided tape, on top of which a small amount of petroleum jelly was spread. ∼100-150 μl of SDM-79 medium was added into the chamber, the glass coverslip, with the cell side facing down, was laid onto the petroleum jelly spread on the double-sided tape. The coverslip was slowly pressed down until it touched the medium. Finally, the chamber was sealed by spreading the nail polish to cover the petroleum jelly.

The slide was used for live-cell imaging with the Nikon A1R Confocal Laser Microscope equipped with a Nikon A1 LFOV camera (aperture: 1.4), using a 60× Apo TIRF/1.49 NA Oil lens. Images were taken every 2 minutes for 15 hours (PFS state: on), using the NIS-Elements AR Imaging Software (v5.41.02, Build 1711, Nikon). Cells were illuminated with laser wavelength of 488 (power: 3.0; PTM HV: 100) for differential interference contrast (DIC) (PTM HV: 140) and 561 (power: 1.8; PTM HV: 67). The pinhole size was set to 62.58 μm, with “Line Average / Integrate Count” of 2, and a Scanner Zoom of 0.72 μm. Images were exported and analyzed using ImageJ.

## Author contributions

**Thiago Souza Onofre:** Conceptualization, Methodology, Visualization, Investigation, Formal analysis, Writing – Reviewing and Editing. **Qing Zhou:** Conceptualization, Methodology, Visualization, Investigation, Formal analysis, Writing – Reviewing and Editing. **Ziyin Li**: Conceptualization, Supervision, Project administration, Funding acquisition, Writing – Original Draft, Writing – Reviewing and Editing.

## Acknowledgements

We thank members of the Li lab for valuable discussions. This work was supported by the NIH grants R01AI118736 and R01AI101437 to Z. L.

## Supplementary data

**Figure S1. Localization of overexpressed, 3HA-tagged TbSPA and its mutants and knock-out of TbSPA in the procyclic form.** (**A**). Immunofluorescence microscopic images of intact cells overexpressing 3HA-tagged TbSPA and its mutants. Scale bar: 5 μm. (**B**). Knock-out of one allele of *TbSPA* gene. Shown are PCR results to confirm the replacement of one allele of *TbSPA* gene with the puromycin-resistance gene.

**Video S1. Time-lapse video of a control cell with normal mitosis.**

**Video S2. Time-lapse video of a TbSPA RNAi with failed mitosis.**

**Video S3. Time-lapse video of a TbSPA RNAi cell with slower mitosis.**

