## Supplementary figures and images for "The microtubule-severing enzyme spastin regulates spindle dynamics to promote chromosome segregation in *Trypanosoma brucei*"

### Figure S1

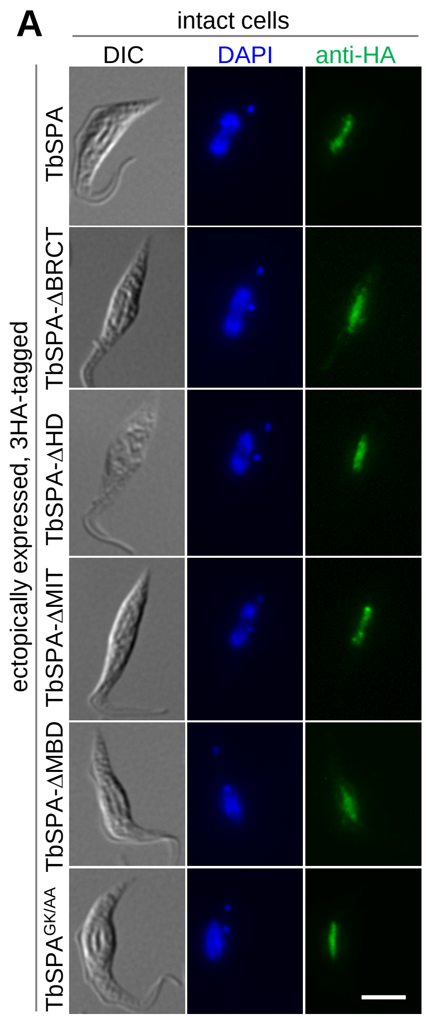

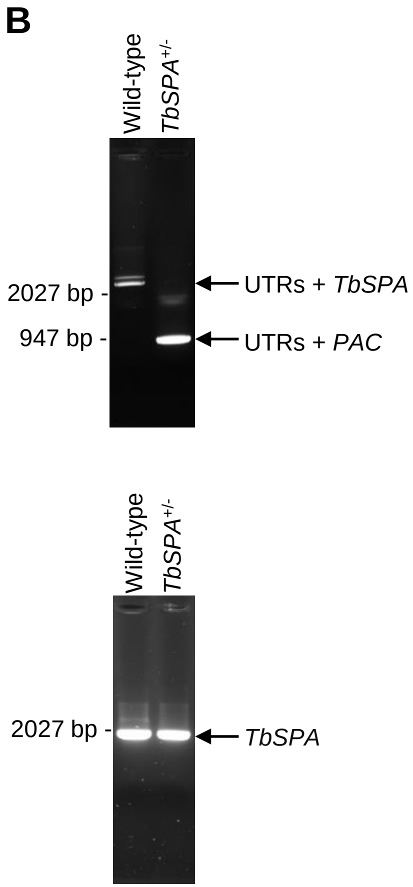
